## Supplementary material for "scE^2^TM improves single-cell embedding interpretability and reveals cellular perturbation signatures": Includes supplementary notes, supplementary figures, and supplementary tables.

### Supplementary materials

**2 Supplementary Figures** **5**

**3 Supplementary Tables** **12**

### 1 Supplementary Notes

#### 1.1 Supplementary Note 1: Details of related work

##### 1.1.1 Single-cell deep embedding methods

With the rapid development of deep learning, deep embedding models have been widely used in the analysis of single cells. Lopez *et al* [1] propose scVI based on the VAE, which approximates the posterior distribution of scRNA-seq data through a variational inference network and reconstructs the cells using a neural decoder. Tian *et al* [2] develop scDCC that combines the Zero-Inflated Negative Binomial (ZINB) model with clustering loss and constraint loss. scDHA proposed by Tran *et al* [3] can utilize a non-negative kernel autoencoder to help remove genes that have little contribution to the representation of the part-based data, thus extracting representative information for each cell more accurately. However, these methods only focus on gene expression information when representing the learning process without considering the topological information of the cell network.

To naturally embed expression and relationship information into potential representations, scGNN [4] utilizes GNNs to integrate structural information of cell-cell networks. GNNs learn the representations of nodes in a graph through neighbor information propagation, taking into account both node features and graph topology. scTAG [5] uses the ZINB model as the backbone and incorporates a topologically adaptive graph convolutional autoencoder to further enhance the modeling capability. scLEGA [6] employs a novel ZINB loss function, which fully takes into account the contribution of genes with lower expression values with combines gene expression and the topological information of cellular network through a multi-point attention mechanism. Nevertheless, the large space occupation of cellular networks and graph convolution algorithms make these models difficult to scale to large single-cell datasets. In addition, these internal knowledge-based guidance paradigms are constrained by the limited information in the given data. Unlike previous approaches, we propose a new paradigm that utilizes the external knowledge obtained from the single-cell foundation model to guide single-cell embedding and clustering analysis. It is worth noting that for a more comprehensive comparison, we use the single-cell foundation model, i.e., scGPT [7], as our ablation.

##### 1.1.2 Single-cell deep interpretable methods

Despite the attractive performance of the deep embedding methods described above, they lack interpretability and fail to provide biologically meaningful characterization of the underlying transcriptome. The study of interpretable deep embedding models is an important direction in scRNA-seq data mining, and a series of models have been developed, which are mainly categorized into a priori knowledge-based and topic-based methods. A priori knowledge-based approach involves injecting a priori biological knowledge into the neural network architecture to improve interpretability. For instance, VEGA [8] enhances VAE with gene annotations whose decoder is a sparse single-layer neural network corresponding to a user-supplied database of gene annotations, thus providing direct interpretability of latent variables. P-NET [9] is a deep learning approach that integrates previously established knowledge of biological hierarchies into a neural network language to guide a supervised learning task, thus making predictions biologically rich and interpretable. TOSICA [10] integrates prior biological knowledge mechanisms through the mechanism of attention. However, these methods are limited by a priori information and suffer from incomplete domain knowledge and difficulty in learning new knowledge.

The single-cell embedded topic model associates topics with genes, thus providing interpretability. scETM model first [11] introduced interpretable linear decoders to learn highly interpretable gene and topic embeddings to extend topic modeling to the single-cell domain. Based on this, SPRUCE [12] combined a topic modeling approach with cell-cell interactions to simulate informational crosstalk between cell-cells while identifying cell types. Morever, Chen *et al* [13] enhance the model's ability to capture latent signals within the data by constructing deeper neural network architectures to better capture the dependencies between topics and genes. Nevertheless, existing topic-based approaches do not provide a quantitative assessment of the interpretability of the model and are not able to cope with potential interpretation collapse.

#### 1.2 Supplementary Note 2: Topic Enrichment Analysis

To assess whether a topic is enriched in any known pathways, a common approach is to detect over-representation analysis (ORA) [14] and Gene Set Enrichment Analysis (GSEA) [15]. ORA contains a background gene set (with known gene functions or pathways) and a specific gene set, and determines whether a specific gene set is enriched for the pathway by identifying the specific gene set as containing a higher proportion of genes than the background gene set, where the specific gene set is selected based on a threshold value. GSEA avoids threshold selection by calculating running sums of enrichment scores down the list of genes, and is one of the commonly used methods for pathway enrichment analysis. There are three key elements of the GSEA method. 1) An enrichment score is computed, which reflects the extent to which the pathway set S is overrepresented at the extremes (top or bottom) of the entire ranked list. 2) Estimate the statistical significance of enrichment score by using an empirical phenotype-based permutation testing program. 3) When evaluating the entire gene set database, we adjusted the estimated significance level to account for multiple hypothesis testing.

In this study, the top-10 genes per topic are used for calculating the topics as well as the metrics for the topic ORA analysis. For the calculation of TCs, we used as external corpus all known gene pathways (msigdb.v2024.1.Hs.symbols and msigdb.v2024.1.Mm.symbols) for mouse and human, respectively, which are downloaded from the GSEA database (`https://www.gsea-msigdb.org/`). Moreover, when performing ORA and GSEA, the external genes we used are curated gene sets from GSEA for humans and mouse.

### 2 Supplementary Figures

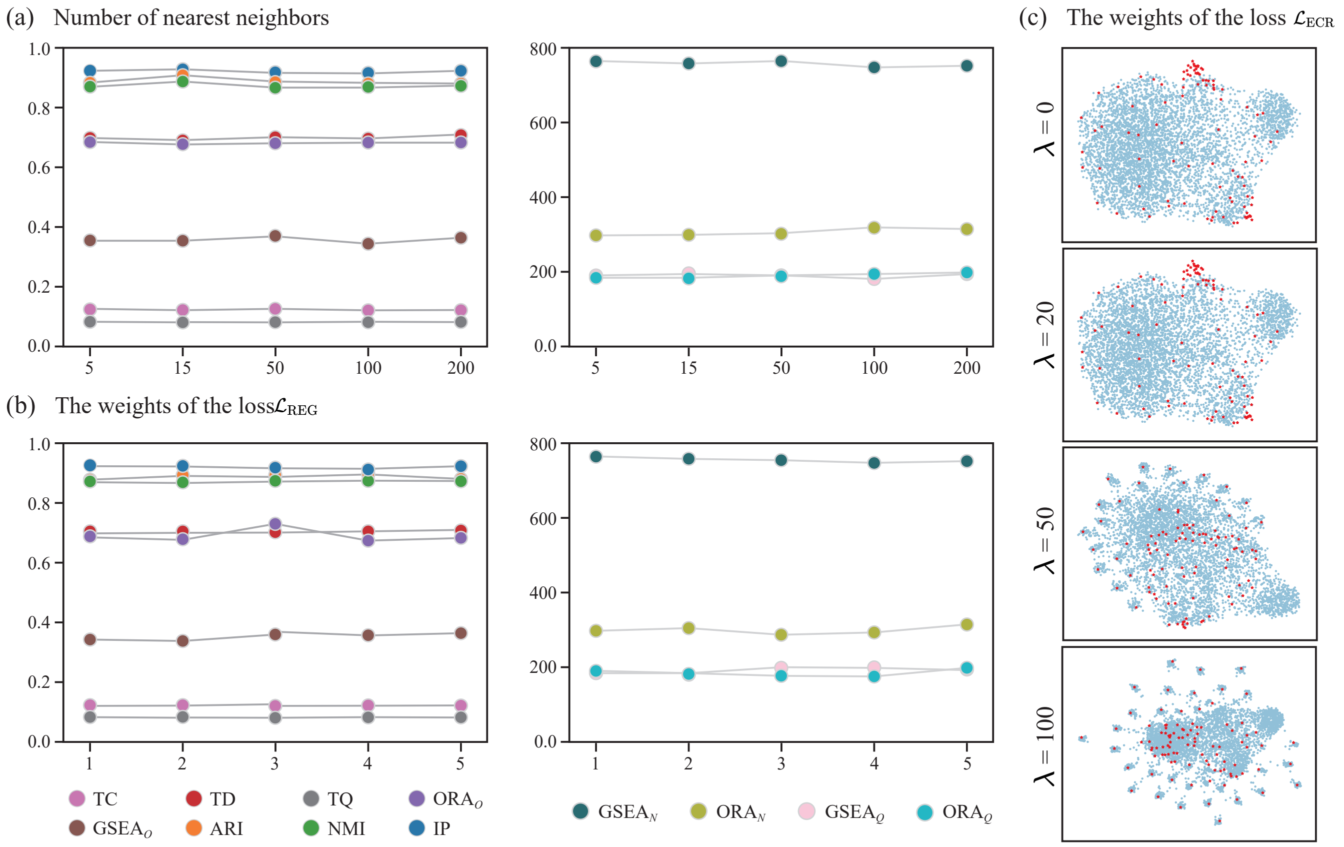

Figure 1: Sensitivity analysis of scE$^2$TM key hyperparameters. (a) Model robustness to neighborhood size in the CVE module. (b) Performance variation with different weights of loss function $\mathcal{L}_{\text{REG}}$. (c) Geometric evolution of topic ($\star$) and gene ($\bullet$) embedding space across $\lambda$ values.

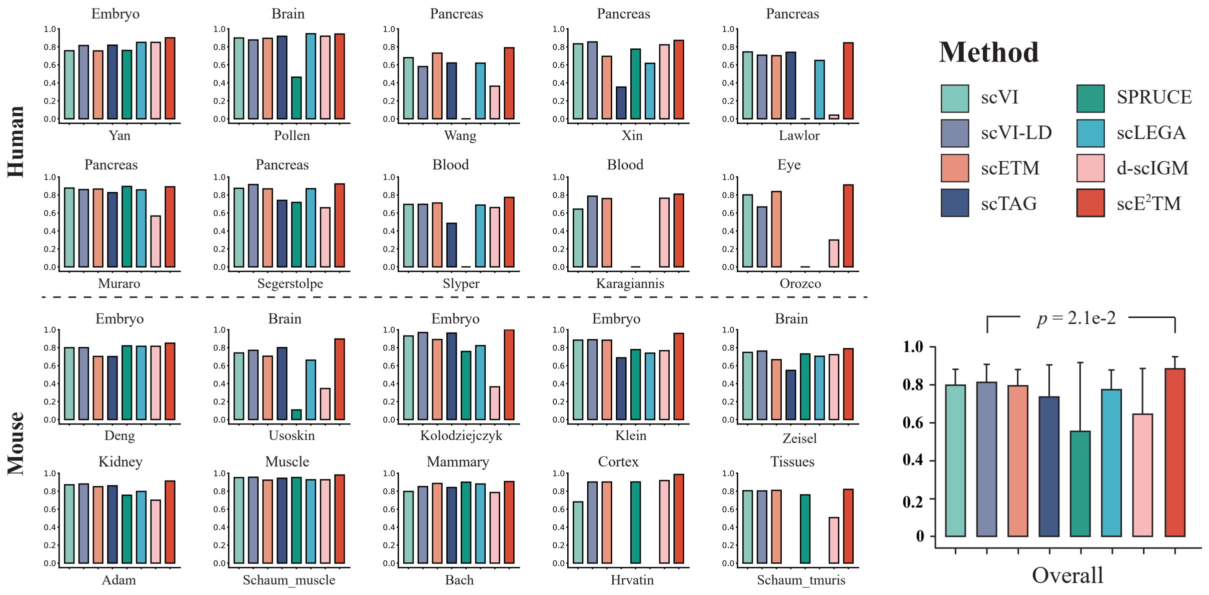

Figure 2: Evaluating clustering performance with NMI. The 20 panels on the left show the NMI values for the different methods on individual dataset, and the right panel shows the average NMI values and standard deviations, with statistical significance tested by pairwise Whitman U-test.

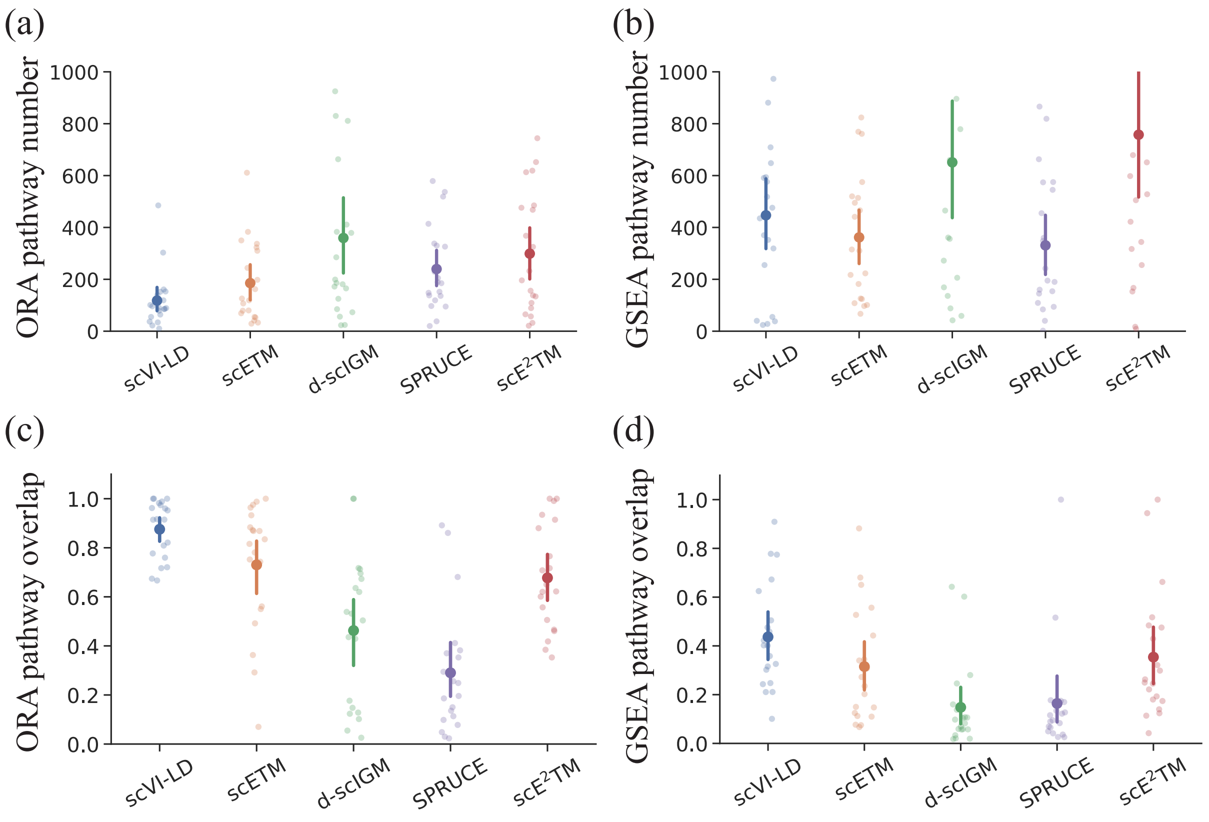

Figure 3: Interpretability evaluation of single-cell embedded topic models.

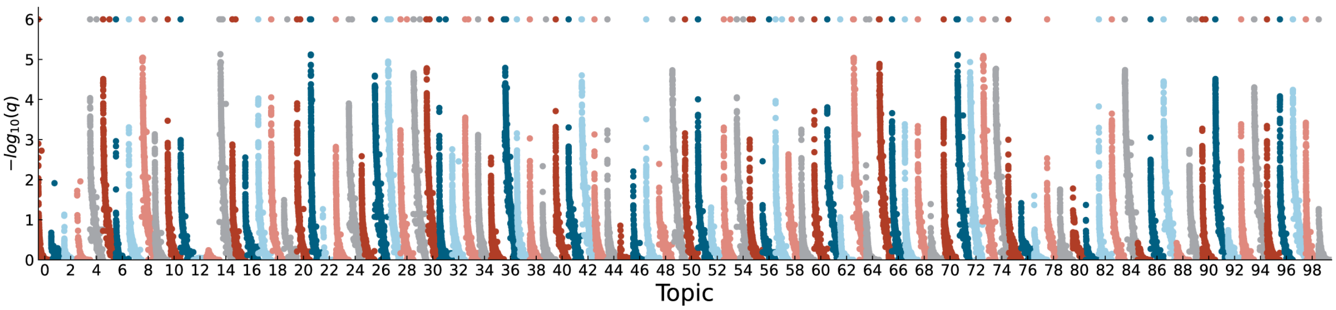

Figure 4: Manhattan plot of GSEA results for 100 topics. The $y$-axis is the significance value from the permutation test corrected.

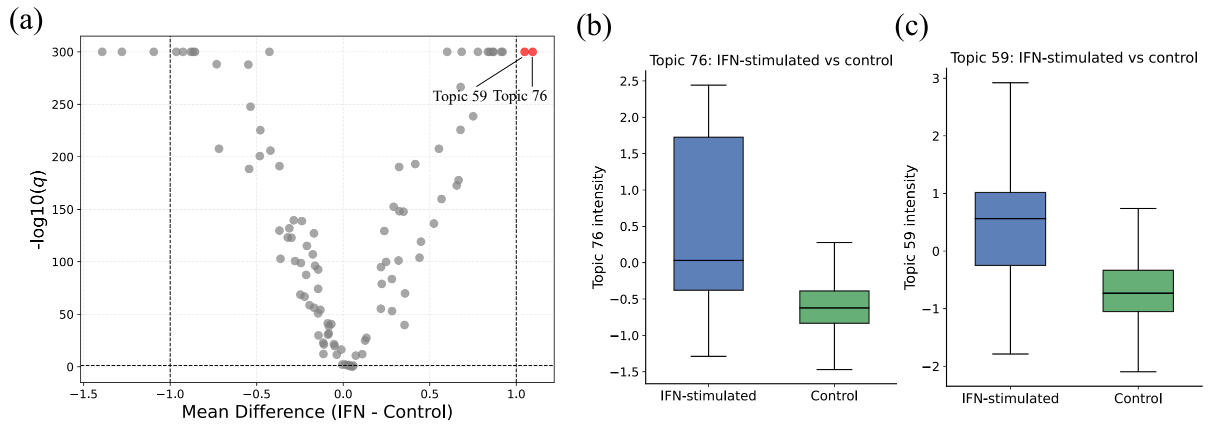

Figure 5: Differential expression analysis of topics on IFN-$\beta$ stimulated datasets. (a) Volcano plot showing differential topic activity between IFN-stimulated and control cells. Topics 76 and 59 (highlighted) exhibit the largest positive mean differences and strongest statistical significance, identifying them as IFN-specific topics. (b) Boxplot of topic 76 activity in IFN-stimulated versus control cells, illustrating a marked increase under IFN stimulation. (c) Boxplot of topic 59 activity, indicating higher activity in IFN-stimulated cells relative to controls.

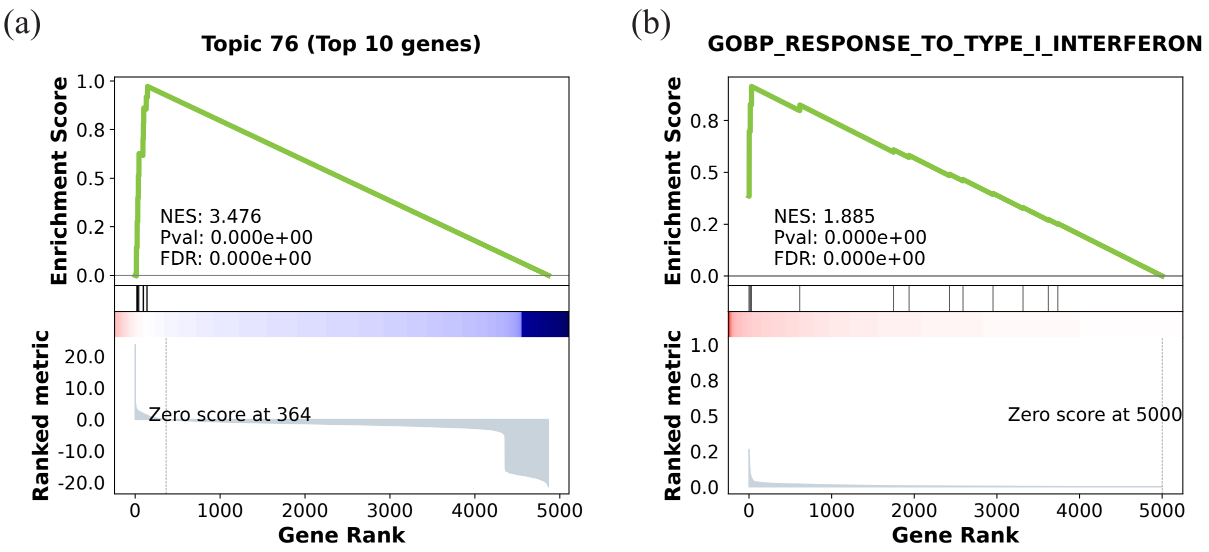

Figure 6: Gene set enrichment analysis. (a) Enrichment analysis between the top 10 genes of topic 76 and the list of differentially expressed genes sorted in descending order by cell type FCGR3A+Mono. (b) Enrichment analysis between the pathway "GOBP_RESPONSE_TO_TYPE_I_INTERFERON" and the descending-sorted gene list of topic 76.

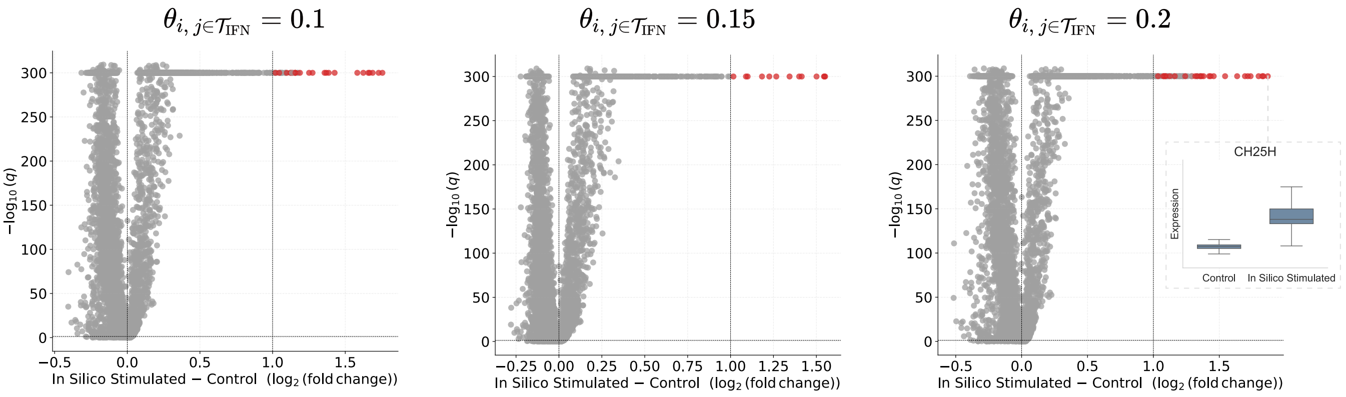

Figure 7: Differential gene expression after in silico stimulation of IFN-specific topics in the PBMC dataset. Volcano plot showing differential gene expression between reconstructed control cells subjected to varying intensities of in silico stimulation and their unstimulated counterparts. The boxplots show the expression differences of the top significantly upregulated genes, CH25H, between perturbed and unperturbed reconstructed cells (IFN-specific topic intensities set to 0.2).

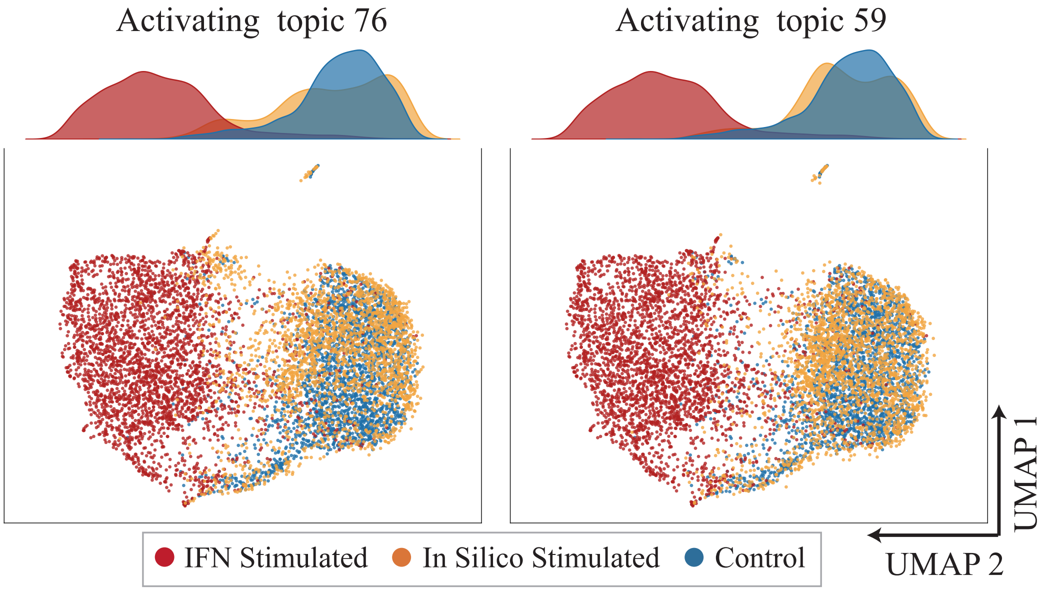

Figure 8: UMAP visualization of cellular responses to perturbing individual IFN-specific topics in the PBMC dataset. Density plots and UMAP embeddings show the distributions of IFN-stimulated, in silico stimulated, and control reconstructed cells after activating IFN-specific topic 76 or topic 59 (IFN-specific topic intensity set to 0.2).

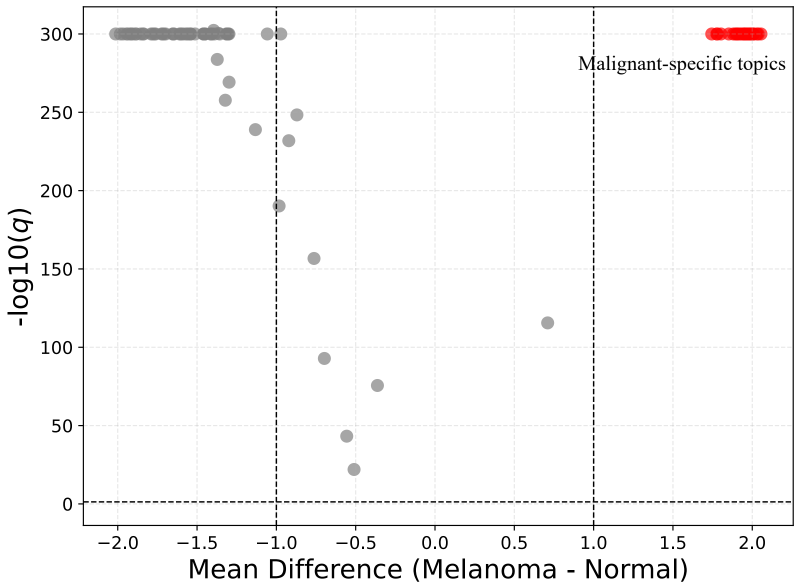

Figure 9: Differential expression analysis of topics on melanoma datasets. Volcano plot showing differential topic activity between malignant and normal cells.

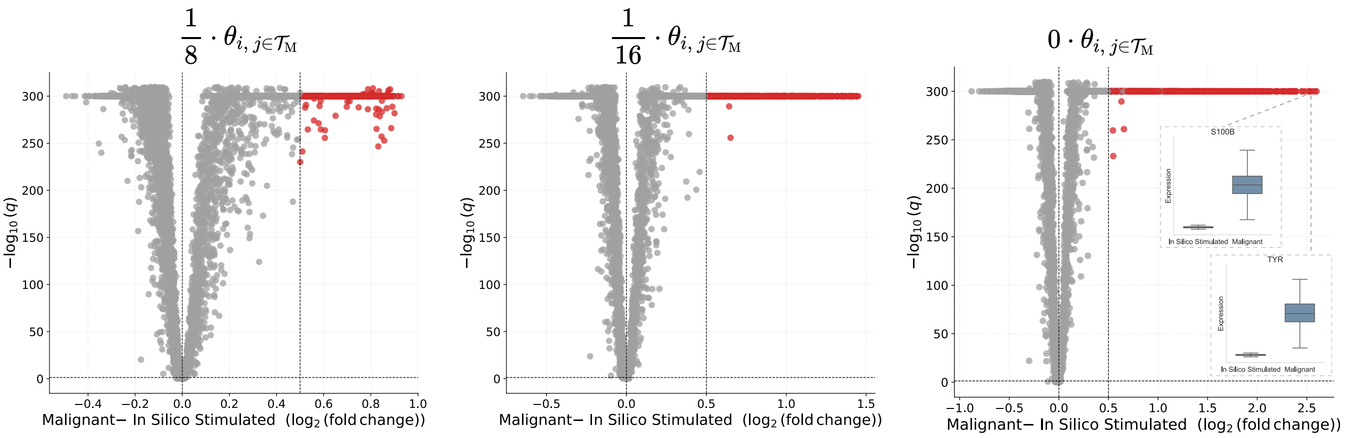

Figure 10: Differential gene expression after in silico stimulation of malignant-specific topics in the melanoma dataset. Volcano plot showing differential gene expression between reconstructed malignant cells subjected to varying intensities of in silico stimulation and their unstimulated counterparts. The boxplots show the expression differences of the top significantly upregulated genes, TYR and S100B, between perturbed and unperturbed reconstructed cells (malignant-specific topic intensities set to 0).

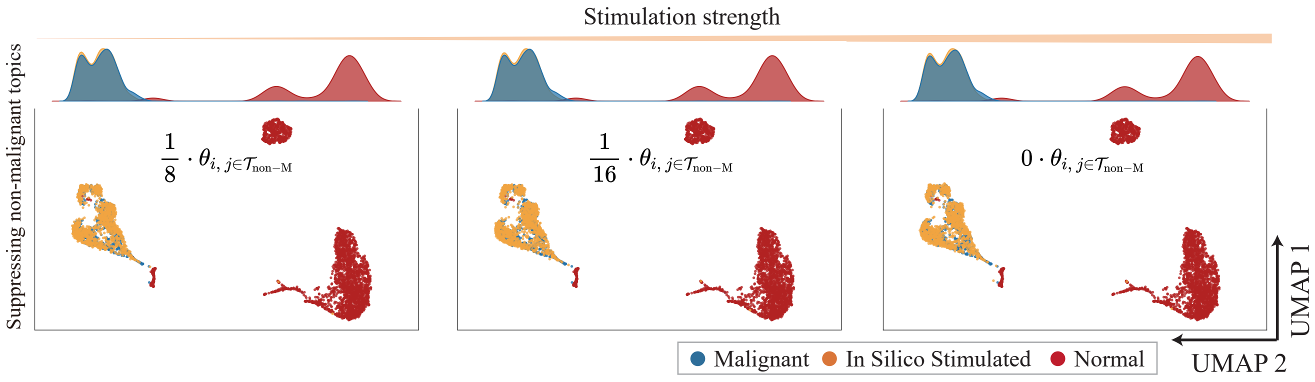

Figure 11: Negative control experiment for the melanoma data. UMAP projections and density maps of reconstructed cells under varying perturbation strength using randomly selected non-melanoma-specific topics as a negative control experiment. $\theta_{i,j}$ denotes the topic intensity of cell $i$ on topic $j$, and $\mathcal{T}_{\text{non-M}}$ denotes the set of non-malignant-specific topics.

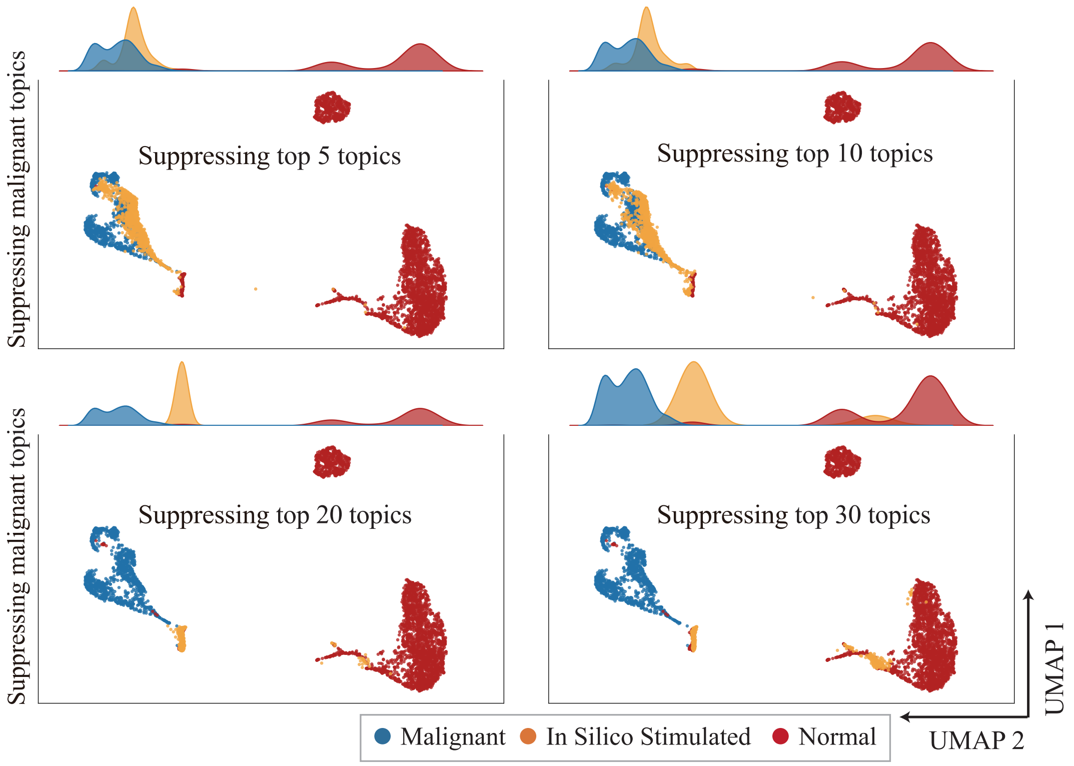

Figure 12: UMAP visualization of cellular responses to perturbing malignant-specific topics in the melanoma dataset. Density plots of cells and UMAP embeddings after suppressing varying numbers of malignant-specific topics. Suppressing the top 5, 10, 20, or 30 malignant-specific topics progressively alters the reconstructed distribution of malignant cells.

### 3 Supplementary Tables

Table 1: Ablation analysis across all datasets. Performance metrics shown as mean ± standard deviation. scGPT produces cell embeddings but not probabilistic topic models, precluding direct comparison on topic interpretability metrics.

| Model | ARI | NMI | TC | TD | TQ | IP | ORA$_U$ | ORA$_N$ | ORA$_Q$ | GSEA$_U$ | GSEA$_N$ | GSEA$_Q$ |
|---|---|---|---|---|---|---|---|---|---|---|---|---|
| scGPT | 0.681$_{\pm 0.23}$ | 0.712$_{\pm 0.18}$ | - | - | - | - | - | - | - | - | - | - |
| scE$^2$TM w/o ECR | 0.862$_{\pm 0.10}$ | 0.862$_{\pm 0.07}$ | **0.147**$_{\pm 0.05}$ | 0.597$_{\pm 0.15}$ | 0.068$_{\pm 0.02}$ | 0.916$_{\pm 0.05}$ | 0.612$_{\pm 0.26}$ | **380**$_{\pm 308}$ | **207**$_{\pm 175}$ | **0.424**$_{\pm 0.25}$ | 688$_{\pm 582}$ | **213**$_{\pm 219}$ |
| scE$^2$TM w/o CVE | 0.843$_{\pm 0.09}$ | 0.845$_{\pm 0.07}$ | 0.119$_{\pm 0.04}$ | 0.700$_{\pm 0.13}$ | 0.080$_{\pm 0.02}$ | 0.913$_{\pm 0.05}$ | **0.694**$_{\pm 0.21}$ | 304$_{\pm 253}$ | 180$_{\pm 160}$ | 0.354$_{\pm 0.27}$ | 750$_{\pm 580}$ | 179$_{\pm 156}$ |
| scE$^2$TM w/ EC | 0.858$_{\pm 0.10}$ | 0.856$_{\pm 0.08}$ | 0.122$_{\pm 0.04}$ | **0.706**$_{\pm 0.14}$ | **0.084**$_{\pm 0.02}$ | 0.929$_{\pm 0.04}$ | 0.683$_{\pm 0.23}$ | 326$_{\pm 256}$ | 198$_{\pm 165}$ | 0.367$_{\pm 0.27}$ | 751$_{\pm 587}$ | 200$_{\pm 230}$ |
| scE$^2$TM | **0.908**$_{\pm 0.08}$ | **0.888**$_{\pm 0.07}$ | 0.122$_{\pm 0.04}$ | 0.690$_{\pm 0.15}$ | 0.081$_{\pm 0.03}$ | **0.930**$_{\pm 0.04}$ | 0.677$_{\pm 0.21}$ | 299$_{\pm 230}$ | 180$_{\pm 142}$ | 0.354$_{\pm 0.257}$ | **758**$_{\pm 586}$ | 195$_{\pm 216}$ |

Table 2: Efficiency analysis of different methods.

| | Wang | | Klein | | Slyper | |
|---|---|---|---|---|---|---|
| **Model** | **Runtime** | **Space** | **Runtime** | **Space** | **Runtime** | **Space** |
| scVI | 40s | 907MB | 195s | 893MB | 842s | 897MB |
| scVI-LD | 37s | 883MB | 187s | 897MB | 729s | 873MB |
| scETM | 140s | 915MB | 760s | 1,167MB | 4,029s | 1,227MB |
| scTAG | 44s | 10,563MB | 106s | 10,563MB | 1,439s | 10,563MB |
| scLEGA | 34s | 1,093MB | 167s | 3,561MB | 244s | 6,051MB |
| SPRUCE | 119s | 739MB | 681s | 761MB | 3,391s | 739MB |
| d-scIGM | 429s | 3,731MB | 2,851s | 3,775MB | 11,154s | 3,777MB |
| scE$^2$TM | 61s | 957MB | 486s | 1,063MB | 2,356s | 1,475MB |

Table 3: The ten most common genes identified in human pancreatic datasets were supported by prior experimental or literature evidence.

| Index | Gene_name | Gene_stable_ID | Reference |
|---|---|---|---|
| 1 | INS | ENSG00000254647 | Andrali *et al* [16] |
| 2 | CEACAM7 | ENSG00000007306 | Raj *et al* [17] |
| 3 | UCA1 | ENSG00000214049 | Chen *et al* [18] |
| 4 | LCN2 | ENSG00000148346 | Gumpper *et al* [19] |
| 5 | PCSK1 | ENSG00000175426 | Wang *et al* [20] |
| 6 | SAA2 | ENSG00000134339 | Chen *et al* [21] |
| 7 | RBP4 | ENSG00000138207 | Fan *et al* [22] |
| 8 | GCG | ENSG00000115263 | Miyazaki *et al* [23] |
| 9 | PENK | ENSG00000181195 | Jin *et al* [24] |
| 10 | NPY | ENSG00000122585 | Myrsen-Axcrona *et al* [25] |

Table 4: Twenty scRNA-seq benchmark datasets

| Dataset | Tissue | #Sample size | #Class | Protocol | Accession ID | Reference |
|---|---|---|---|---|---|---|
| Yan | Human embryo | 90 | 6 | Tang | GSE36552 | Yan *et al* [26] |
| Deng | Mouse embryo | 268 | 6 | Smart-Seq2 | GSE45719 | Deng *et al* [27] |
| Pollen | Human brain | 301 | 11 | SMARTer | SRP041736 | Pollen *et al* [28] |
| Wang | Human pancreas | 457 | 7 | SMARTer | GSE83139 | Wang *et al* [29] |
| Lawlor | Human pancreas | 604 | 9 | SMARTer | GSE86473 | Lawlor *et al* [30] |
| Usoskin | Mouse brain | 622 | 4 | STRT-Seq | GSE59739 | Usoskin *et al* [31] |
| Kolodziejczyk | Mouse embryo stem cells | 704 | 3 | SMARTer | E-MTAB-2600 | Kolodziejczyk *et al* [32] |
| Xin | Human pancreas | 1,600 | 8 | SMARTer | GSE81608 | Xin *et al* [33] |
| Muraro | Human pancreas | 2,126 | 10 | CEL-Seq2 | GSE85241 | Muraro *et al* [34] |
| Segerstolpe | Human pancreas | 2,209 | 14 | Smart-Seq2 | E-MTAB-5061 | Segerstolpe *et al* [35] |
| Klein | Mouse embryo stem cells | 2,717 | 4 | inDrop | GSE65525 | Klein *et al* [36] |
| Zeisel | Mouse brain | 3,005 | 9 | STRT-Seq | GSE60361 | Zeisel *et al* [37] |
| Adam | Mouse kidney | 3,660 | 8 | Drop-seq | GSE94333 | Adam *et al* [38] |
| Schaum_muscle | Mouse limb muscle | 3,909 | 6 | 10X Genomics | GSE109774 | Schaum *et al* [39] |
| Slyper | Human blood | 13,316 | 8 | 10X Genomics | SCP345 | Single Cell Portal [40] |
| Bach | Mouse mammary epithelial cells | 23,184 | 8 | 10X Genomics | GSE106273 | Bach *et al* [41] |
| Hrvatin | Mouse visual cortex | 48,266 | 8 | inDrop | GSE102827 | Hrvatin *et al* [42] |
| Schaum_tmuris | Mouse tissues | 54,439 | 40 | 10X genomics | GSE109774 | Schaum *et al* [39] |
| Karagiannis | Human blood | 72,914 | 12 | 10X genomics | GSE128879 | Karagiannis *et al* [43] |
| Orozco | Human eye | 100,055 | 11 | 10X genomics | GSE135133 | Orozco *et al* [44] |

40. Single cell portal. https://singlecell.broadinstitute.org/single_cell/study/scp345/.

41. Karsten Bach, Sara Pensa, Marta Grzelak, James Hadfield, David J Adams, John C Marioni, and Walid T Khaled. Differentiation dynamics of mammary epithelial cells revealed by single-cell rna sequencing. *Nature Communications*, 8(1):1–11, 2017.

42. Sinisa Hrvatin, Daniel R Hochbaum, M Aurel Nagy, Marcelo Cicconet, Keiramarie Robertson, Lucas Cheadle, Rapolas Zilionis, Alex Ratner, Rebeca Borges-Monroy, Allon M Klein, et al. Single-cell analysis of experience-dependent transcriptomic states in the mouse visual cortex. *Nature Neuroscience*, 21(1):120–129, 2018.

43. Tanya T Karagiannis, John P Cleary Jr, Busra Gok, Andrew J Henderson, Nicholas G Martin, Masanao Yajima, Elliot C Nelson, and Christine S Cheng. Single cell transcriptomics reveals opioid usage evokes widespread suppression of antiviral gene program. *Nature Communications*, 11(1):2611, 2020.

44. Luz D Orozco, Hsu-Hsin Chen, Christian Cox, Kenneth J Katschke, Rommel Arceo, Carmina Espiritu, Patrick Caplazi, Sarajane Saturnio Nghiem, Ying-Jiun Chen, Zora Modrusan, et al. Integration of eqtl and a single-cell atlas in the human eye identifies causal genes for age-related macular degeneration. *Cell reports*, 30(4):1246–1259, 2020.
